# Homeostatic activation of Aryl Hydrocarbon Receptor by dietary ligands dampens cutaneous allergic responses by controlling Langerhans cells migration

**DOI:** 10.1101/2023.01.24.525336

**Authors:** Adeline Cros, Alba de Juan, Renaud Leclere, Julio L Sampaio, Mathieu Maurin, Sandrine Heurtebise-Chrétien, Elodie Segura

**Author notes:** **Correspondence:** Elodie Segura.

## Abstract

Dietary compounds can affect the development of inflammatory responses at distant sites. However, mechanisms involved remain incompletely understood. Here we addressed the influence on allergic responses of dietary agonists of Aryl Hydrocarbon Receptor (AhR). In cutaneous papain-induced allergy, we found that lack of dietary AhR ligands exacerbates allergic responses. This phenomenon was tissue-specific, as airway allergy was unaffected by the diet. In addition, lack of dietary AhR ligands worsened asthma-like allergy in a model of ‘atopic march’. Mice deprived of dietary AhR ligands displayed impaired Langerhans cell migration, leading to exaggerated T cell responses. Mechanistically, dietary AhR ligands regulated the inflammatory profile of epidermal cells, without affecting barrier function. In particular, we evidenced TGF-β hyperproduction in the skin of mice deprived of dietary AhR ligands, explaining Langerhans cell retention. Our work identifies an essential role for homeostatic activation of AhR by dietary ligands in the dampening of cutaneous allergic responses and uncovers the importance of the gut-skin axis in the development of allergic diseases.

## Introduction

Development of immune-mediated diseases is affected by numerous environmental factors, including nutrition. Dietary compounds can modulate immune cells homeostasis and inflammatory immune responses (Wu et al., 2019), affecting in particular the susceptibility to allergy (Julia et al., 2015). However, the impact of individual nutrients and the molecular mechanisms involved remain incompletely understood. In particular, whether dietary metabolites that activate the Aryl Hydrocarbon receptor (AhR) influence type 2 allergic responses remains unclear.

Type 2 allergic responses result from dysregulated immune responses mediated mainly by IL4, IL5 and IL13. In the skin and lungs, allergens trigger the secretion by keratinocytes or epithelial cells of alarmins, such as thymic stromal lymphopoietin (TSLP) (Deckers et al., 2017a; Akdis et al., 2020). These soluble mediators stimulate the production of IL4, IL5 and IL13 by type 2 innate-like lymphoid cells (ILC) and basophils, and the activation of dendritic cells (DC) (Deckers et al., 2017a; Akdis et al., 2020). After their migration to the lymph nodes, activated DC polarize CD4 T cells into Th2 cells which themselves secrete IL4, IL5 and IL13, thereby stimulating the production of IgE and amplifying the type 2 inflammatory response (Deckers et al., 2017a; Akdis et al., 2020). Nutritional compounds can affect multiple players of the type 2 inflammatory cascade through recognition by G-protein coupled and nuclear receptors. For instance, mice fed on a high-fiber diet have increased levels of short-chain fatty acid propionate and decreased susceptibility to airway allergy, due to impaired induction of Th2 polarization by lung DC (Trompette et al., 2014). Mice with a Vitamin D-deficient diet display lung inflammation, high blood IgE levels and increased numbers of lung Th2 cells (Vasiliou et al., 2014). Dietary interventions are therefore a promising strategy for preventing the development of allergic diseases, in particular in children (Trambusti et al., 2020; Trikamjee et al., 2021). However, a better understanding of the effects of each family of dietary compounds is needed.

AhR is a nuclear receptor sensing metabolites produced mainly by the breakdown of food components or from tryptophan catabolism by intestinal microbiota (Rothhammer and Quintana, 2019; De Juan and Segura, 2021). Imbalance in gut-derived AhR ligands worsens inflammation in the intestine during inflammatory bowel disease (Monteleone et al., 2011; Lamas et al., 2016; Hubbard et al., 2017) and in the central nervous system during neuro-inflammation (Rothhammer et al., 2018, 2016). Whether dietary AhR ligands also play a role in other pathological contexts such as allergy requires further investigation.

In this study, we explored the role of dietary AhR ligands in the development of type 2 allergic responses using papain as a model allergen. We showed that lack of dietary AhR ligands exacerbates cutaneous, but not airway, papain-induced allergy. In addition, we found that lack of dietary AhR ligands during allergen epicutaneous sensitization worsened asthma-like airway allergy in a recall phase, a model for ‘atopic march’. We demonstrated that lack of dietary AhR ligands impacts the inflammatory profile of epidermal cells and increases the production of bioactive TGF-β, causing the retention of Langerhans cells in the skin, in turn leading to exaggerated Th2 responses in the lymph nodes. Our results identify a major role for dietary AhR ligands in the modulation of cutaneous allergic responses.

## Results

### Lack of dietary AhR ligands exacerbates cutaneous allergic type 2 responses

To study the impact of dietary AhR agonists on allergic responses, we sought to compare mice fed with diets either poor or rich in AhR agonists. Normal mouse chow contains phytochemicals that can act as precursors of AhR agonists, in particular indole-3-carbinole (I3C) (Bjeldanes et al., 1991) which is detected in the serum of mice fed on normal chow diet (FigS1A). To avoid confounding effects, we used a standard purified diet (AIN-93M) which is naturally poor in phytochemicals (hereafter termed ‘AhR-poor diet’) and the same diet enriched for I3C (hereafter termed ‘I3C diet’). As a proof-of-principle, we verified that I3C concentration in the serum of mice fed on the AhR-poor or I3C diet was significantly different (FigS1A). To confirm that these diets induced different levels of AhR activation *in vivo*, we analyzed the expression in liver cells of the canonical AhR target gene *Cyp1a1*. Mice fed on the normal chow and I3C diets showed similar expression of *Cyp1a1*, while mice fed on the AhR-poor diet had almost undetectable expression of *Cyp1a1* (Fig.S1B), validating that the AhR-poor diet contains negligeable amounts of natural AhR agonists. To study type 2 allergic responses, we chose the model protease allergen papain (Shimura et al., 2016; Kamijo et al., 2013; Sokol et al., 2008). To validate that the I3C diet would mimick normal conditions in this model, we analyzed Th2 cells induction in mice fed on normal chow or I3C diet. After footpad injection of papain, we observed similar Th2 induction in the draining lymph nodes in both groups (Fig.S1C). IL4 and IL13 were increased by papain exposure, while IL5 secretion was not significantly different from the control condition. Finally, to confirm that only AhR agonists from the diet were modulated in this setting, we measured the serum concentration of L-kynurenin (produced by host metabolism), 3,3’-Diindolylmethane (DIM, generated by the degradation of I3C in the stomach), and Indole-3-acetic acid (produced by microbiota metabolism from food components) (De Juan and Segura, 2021; Bjeldanes et al., 1991). DIM, but not L-kynurenin or Indole-3-acetic acid, was decreased in the serum of mice fed on the AhR-poor diet (FigS1D). Collectively, these results validate our experimental set up. For the rest of the study, we compared mice fed on the I3C and AhR-poor diets.

We first analyzed cutaneous allergic responses to papain challenge in the footpad. Histological analysis showed epidermal hyperplasia upon papain exposure, only in the foot of mice fed on the AhR-poor diet (Fig.1A-B and FigS1E). To assess the induction of Th2 responses, we analyzed draining lymph nodes after 6 days. The number of CD4 T cells and B cells was similar in mice fed on the IC3 or AhR-poor diets (Fig.1C), suggesting that the diet had no impact on CD4 T cells or B cells proliferation. We restimulated normalized numbers of lymph nodes T cells *ex vivo*,and measured cytokine secretion. While IL4 secretion was induced similarly in both groups of mice, the production by T cells of Th2 cytokines IL5 and IL13 was exacerbated in mice fed on the AhR-poor diet (Fig.1C), as well as the secretion of IL10, IFN-γ and IL17A (Fig.S1F). We concluded that lack of dietary AhR ligands amplifies allergic responses after cutaneous allergen exposure.

**Figure 1.**
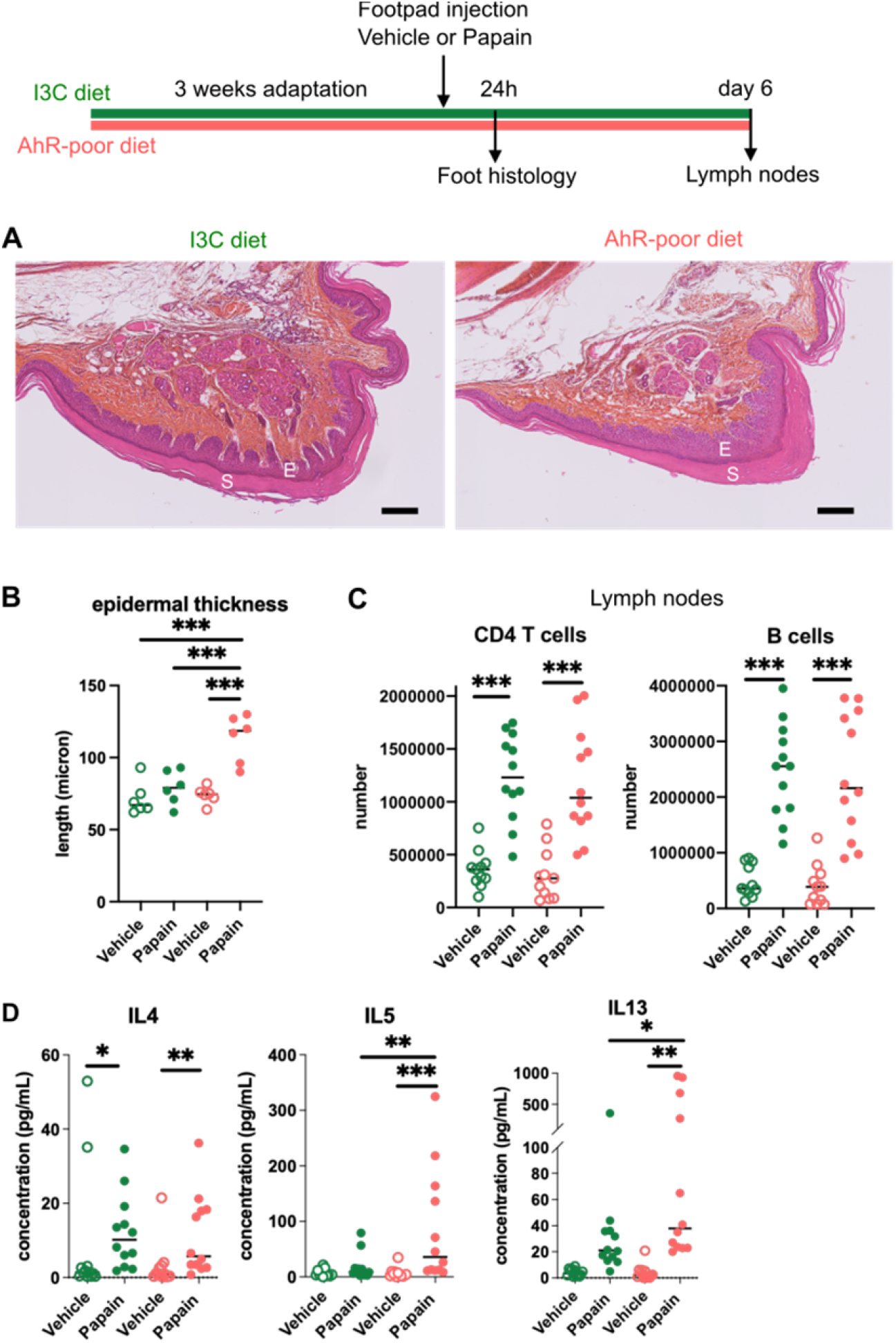
Lack of dietary AhR ligands exacerbates cutaneous allergic type 2 responses. Mice were fed on AhR-poor diet or enriched in Indole-3-carbinol (I3C diet). Papain or vehicle (PBS) was injected in the footpad. (A-B) Tissues were analyzed by histology 24h after papain injection (Hematoxylin, Eosin and Safran staining). (A) Representative results (n=6 per condition). E=epidermis, *S=stratum corneum*. Bar = 100 μm. (B) Epidermal thickness was measured on images. Median is shown (n=6 in 2 independent experiments). One-way ANOVA. (C) After 6 days, cells from the draining lymph nodes were analyzed. (C) CD4 T cells and B cells counts. (D) Normalized numbers of T cells were restimulated *ex vivo* and cytokine secretion was measured in the supernatant after 24h. Median is shown (n=11-12 in 3 independent experiments). Kruskal-Wallis test. For all panels * p<0.05; ** p<0.01; *** p<0.001.

### Lack of dietary AhR ligands does not impact airway allergic type 2 inflammation

To address whether this phenomenon occurred with other exposure routes, we used a model of asthma-like airway allergy induced by repetitive intra-nasal exposure to papain. Papain challenge induced airway inflammation with infiltration of eosinophils, monocytes and CD4 T cells as assessed in the broncho-alveolar space (Fig.2A and Fig.S2). There was no difference in cell numbers between mice fed on the I3C or AhR-poor diets (Fig.2A). We could also detect IL13 and IL5 in the broncho-alveolar lavage, but cytokine secretion was not increased by the AhR-poor diet compared to I3C diet (Fig.2B). The production of Th2 cytokines by T cells from the pulmonary lymph nodes was also similar between diets (Fig.2C). We also assessed airway hyperreactivity by measuring in lung tissues the expression of the genes coding for mucus protein Mucin 5a (*Muc5ac*) and CLCA1 (*Gob5*), a molecule produced by goblet cells during hyperplasia (Patel et al., 2009). Papain challenge increased the expression of *Muc5ac* and *Gob5* in the lungs, to a similar extent in mice fed on I3C or AhR-poor diets (Fig2D). Finally, we measured the concentration of IgE in the blood, and found comparable levels in both groups of papain-exposed mice (Fig.2E). Collectively, these results indicate that the lack of dietary AhR ligands does not affect airway type 2 allergic responses.

**Figure 2.**
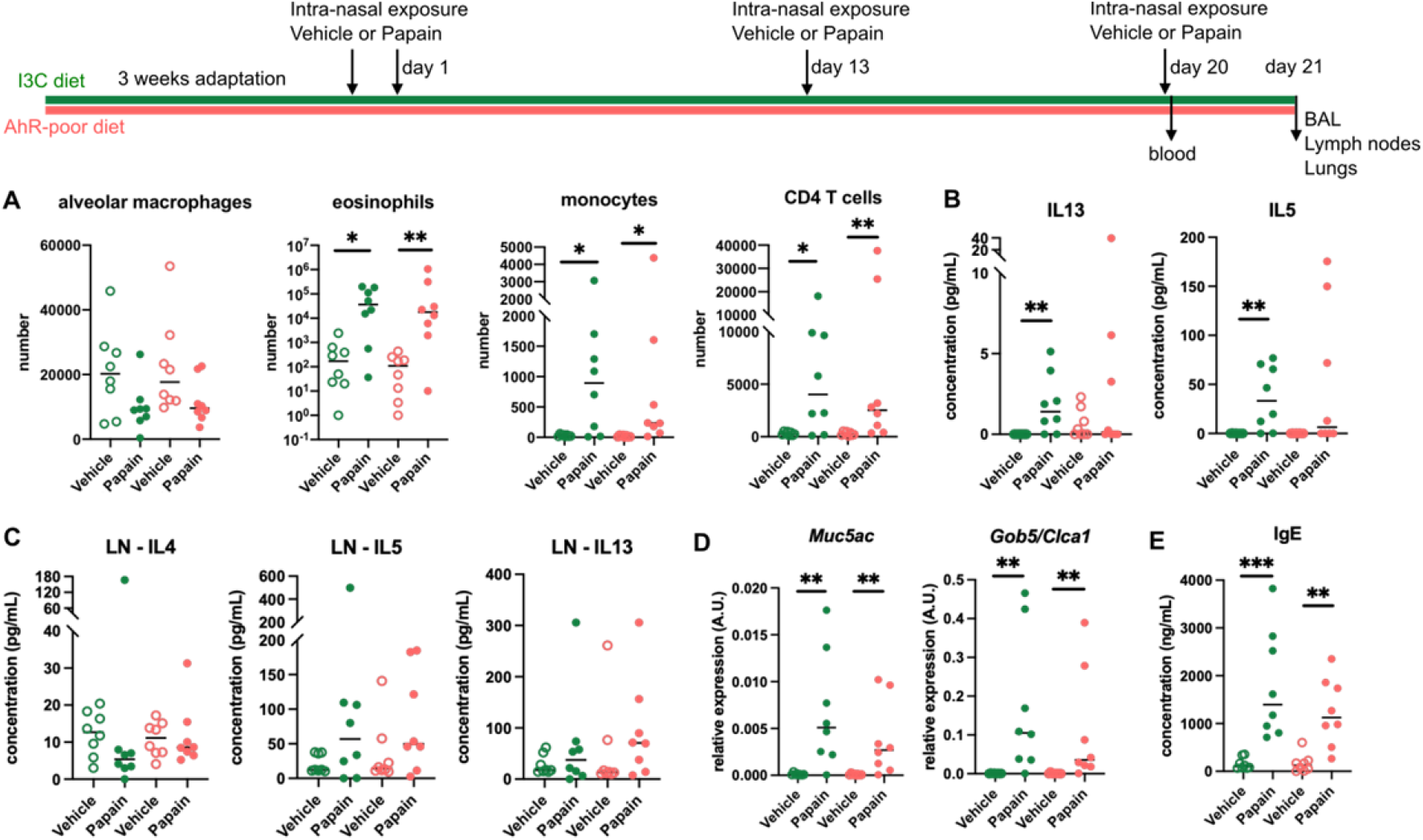
Lack of dietary AhR ligands does not impact airway allergic type 2 inflammation. Mice were fed on AhR-poor or I3C diet. Mice were exposed to papain or vehicle (PBS) intra-nasally 4 times at day 0, 1, 13 and 20. (A-D) 24h after the last exposure, tissues were analyzed. (A) Cell counts in broncho-alveolar lavage. (B) Cytokine concentration in broncho-alveolar lavage. (C) Normalized numbers of lymph nodes T cells were restimulated ex vivo and cytokine secretion was measured in the supernatant after 24h. LN=lymph nodes. (D) Lung lysates were analyzed by RT-qPCR. (E) Blood was collected at day 20 and IgE concentration measured in the serum. For all panels, median is shown (n=8 in 2 independent experiments). Kruskal-Wallis test. * p<0.05; ** p<0.01; *** p<0.001.

### Lack of dietary AhR ligands worsens airway allergy after skin sensitization

Epicutaneous allergen exposure can lead to airway allergy to the same allergen, via the induction of Th2 memory CD4 T cells by skin DC (Deckers et al., 2017a), a phenomenon referred to as ‘atopic march’ (Bantz et al., 2014). Therefore we addressed whether dietary AhR ligands impact asthma-like airway allergy after skin sensitization. To avoid confounding factors, we limited the diet intervention to the sensitization phase, and switched all experimental groups to I3C diet 7 days after cutaneous exposure to papain or vehicle. Then, all groups were exposed to papain intra-nasally. While the number of alveolar macrophages was unaffected by the diet, lack of dietary AhR ligands during the sensitization phase increased the infiltration in the broncho-alveolar space of eosinophils, monocytes and CD4 T cells (Fig.3A). IL13 secretion was higher in the broncho-alveolar lavage of mice fed on the AhR-poor diet during the sensitization phase, and IL5 concentration showed an increased tendancy that was not statistically significant (Fig.3B). Finally, the expression of *Gob5* in lung tissues was significantly increased in mice deprived of dietary AhR ligands during the sensitization phase (Fig.3C). These results show that the lack of dietary AhR ligands during epicutaneous allergen exposure worsens subsequent airway allergy.

**Figure 3.**
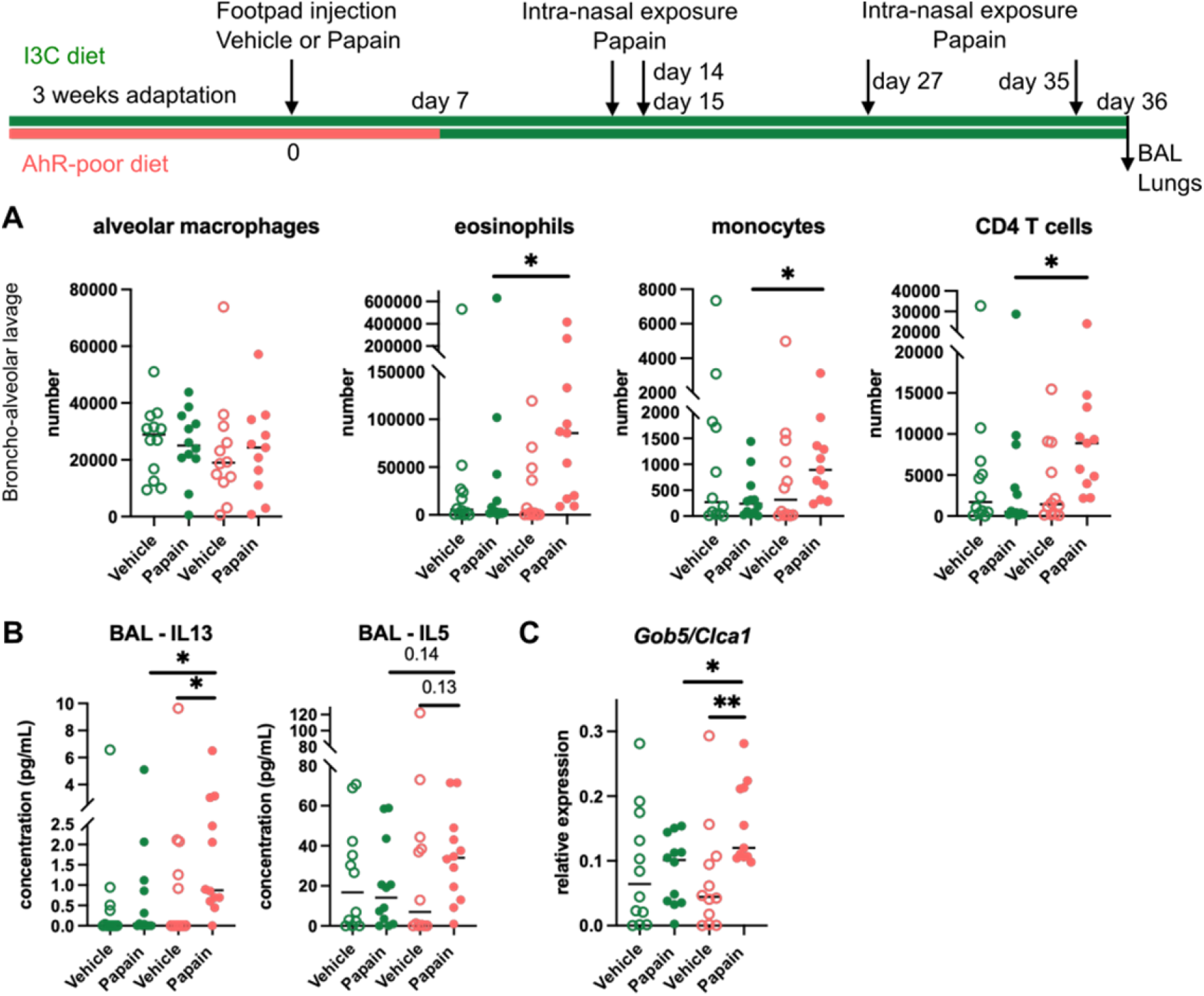
Lack of dietary AhR ligands worsens airway allergy after skin sensitization. Mice were fed on AhR-poor or I3C diet. Papain or vehicle (PBS) was injected in the footpad at day 0. At day 7, all mice were placed on the I3C diet. Mice were exposed to papain intra-nasally 4 times at day 14, 15, 27 and 35. 24h after the last exposure, broncho-alveolar lavage (BAL) and lung tissue were analyzed. (A) Cell counts in BAL. (B) Cytokine concentration in BAL. (C) Lung lysates were analyzed by RT-qPCR. For all panels, median is shown (n=11-12 in 3 independent experiments). Kruskal-Wallis test. * p<0.05; ** p<0.01.

### Lack of dietary AhR ligands impairs Langerhans cells migration which increases Th2 responses in the lymph nodes

The contrasting results obtained in cutaneous versus airway papain-induced allergy suggest tissue-specific effects. Because of their central role in Th2 cells induction, we analyzed DC populations. One difference between skin and lung tissues is the presence of Langerhans cells, residing in the epidermis and absent from the lungs. AhR is expressed by Langerhans cells and has been proposed to regulate their numbers in the epidermis (Hong et al., 2020). To address whether lack of dietary AhR ligands affects Langerhans cells maintenance in the epidermis, we quantified epidermal Langerhans cells from mice fed on the I3C or AhR-poor diet. We first imaged the epidermis of mice expressing Green Fluorescent Protein (GFP) under the promoter of the *Langerin* gene (Kissenpfennig et al., 2005) (Fig.4A). We found no significant difference between diets in Langerhans cells density. To confirm this observation using another approach, we analyzed footpad epidermal cells by flow cytometry (Fig.S3A-B), and found no significant difference between diets in Langerhans cells numbers. We then analyzed skin DC migration to the skin-draining lymph nodes. We assessed DC numbers 24h and 48h after papain injection, compared with vehicle (Fig.S3C-D). Langerhans cells numbers in the lymph nodes was significantly lower upon papain exposure in mice fed on the AhR-poor diet (Fig.4B). By contrast, papain exposure increased the number of migratory skin cDC1 and cDC2 independently of the diet (Fig.4B). In particular, migratory PDL2^+^ cDC2 play an essential role in the induction of allergic Th2 responses (Gao et al., 2013; Kumamoto et al., 2013), and their number was similar in papain-treated mice from both diet groups (Fig.4B). In addition, the number of lymph node resident DC was similar in both diet groups (Fig.4B). The expression of MHC class II molecules or costimulatory molecule CD40 was not affected by the diet (FigS3E). Collectively, our results show that lack of dietary AhR ligands impairs Langerhans cells migration upon papain challenge.

**Figure 4.**
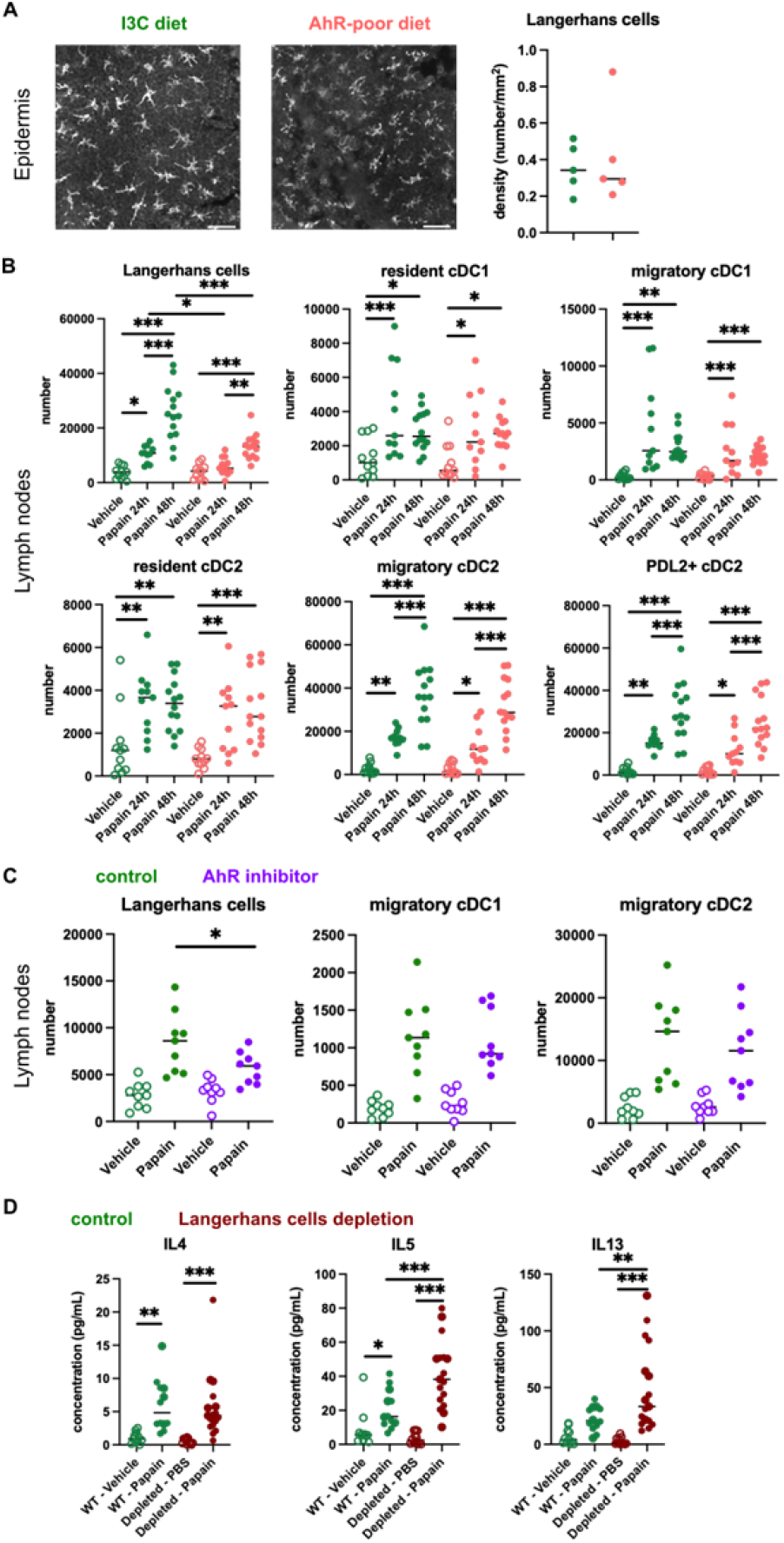
Lack of dietary AhR ligands impairs Langerhans cells migration thereby increasing Th2 responses. (A-B) Mice were fed on AhR-poor or I3C diet. (A) Density of epidermal Langerhans cells was assessed by imaging *Langerin*-eGFP+ cells. Representative images are shown (n=5 per condition). Median is shown. (B-D) Papain or vehicle (PBS) was injected in the footpad. (B) After 24h (vehicle and Papain) or 48h (Papain), dendritic cells numbers were assessed in the draining lymph nodes. Median is shown (n=11-14 in 3 independent experiments). (CD) Mice were fed with the I3C diet. (C) Mice were treated with vehicle or AhR inhibitor CH-223191. After 48h, dendritic cells numbers were assessed in the draining lymph nodes. Median is shown (n=9 in 3 independent experiments). (D) *Langerin-DTR* mice or WT littermates were injected with Diphteria Toxin 3 days prior to Papain treatment. After 6 days, cells from the draining lymph nodes were analyzed. Normalized numbers of lymph nodes T cells were restimulated ex vivo and cytokine secretion was measured in the supernatant after 24h. Median is shown (n=11-17 in 3 independent experiments). One-way ANOVA. For all panels * p<0.05; ** p<0.01; *** p<0.001.

To confirm the role of AhR in this phenomenon, we examined DC migration after papain challenge upon pharmacological inhibition of AhR. We used CH-223191, a potent AhR antagonist (Zhao et al., 2010), to treat mice fed on the I3C diet. Langerhans cells migration, but not that of cDC1 or cDC2, was significantly reduced in mice treated with the AhR inhibitor (Fig.4C). This result suggests that impaired Langerhans cells migration in mice fed with the AhR-poor diet is due to reduced AhR activation.

Langerhans cells play an essential role in dampening T cell responses in skin-draining lymph nodes (Igyarto et al., 2009; Kaplan et al., 2005; Gomez de Agüero et al., 2012). We hypothesized that reduced Langerhans cells presence in the lymph nodes could cause the observed exacerbated Th2 responses. To address this, we used *Langerin-DTR* mice, in which Langerhans cells can be depleted by Diphteria Toxin injection (Kissenpfennig et al., 2005). We confirmed efficient Langerhans cells depletion in the footpad epidermis (Fig.S3F). We injected papain or vehicle in the footpad of mice depleted of Langerhans cells or WT littermates (treated similarly with Diphteria Toxin). To assess Th2 cells induction, we analyzed draining lymph nodes after 6 days. While IL4 secretion was similar in both groups, IL5 and IL13 secretion was significantly increased by Langerhans cells depletion (Fig.4D). These results mirror the observations made in mice fed with the AhR-poor diet (Fig.1D). We concluded that exacerbated Th2 responses in the absence of dietary AhR ligands are caused by impaired Langerhans cells migration to the lymph nodes.

### Dietary AhR ligands regulate the inflammatory profile of epidermal cells

To address how dietary AhR ligands control Langerhans cells migration, we analyzed the transcriptomic profile of epidermal cells in mice fed on the I3C or AhR-poor diet, in basal conditions (vehicle treatment) or upon allergen challenge (6h after papain treatment) (Table S1). Papain exposure induced a common transcriptomic program (Fig.S4A), enriched for type I interferon pathway, cytokine-mediated signaling, matrix remodelling and oncostatin M pathway (a regulator of keratinocyte activation) (Boniface et al., 2007) (Fig.S4B). Consistent with pathway enrichment results, both papain-treated groups had significantly increased expression of inflammatory mediators genes (such as *Csf1, Ccl20* and *Tslp*), interferon-stimulated genes (including *Stat1* and *Oasl2*) (Fig.S4C) and genes involved in tissue repair and remodelling (such *as Areg, Il24, Mmp13, Osmr, Tgfa* and *Vegfa*) (Fig.S4D).

Differentially expressed genes between diets were over-expressed mostly in the epidermis of mice fed on the AhR-poor diet (Fig.5A), both in basal conditions and upon papain challenge (Fig.S4E). Diet-modulated genes were enriched for regulation of extracellular matrix, cytokine signaling and inflammatory responses including the leptin pathway (Fig.5B-C). Consistent with this, mice fed on the AhR-poor diet had in papain-treated epidermis significantly higher expression of matrix components (such as *Col1a1, Col4a1, Col5a1*) and cell adhesion molecules (including *Itgb2* and *Itgb7*) (Fig.S4F), and inflammatory genes including cytokine *Il1b* (Fig.5D) and chemokines *Cxcl1, Cxcl3, Cxcl5, Ccl2* and *Ccl3* (Fig.5D). To confirm these results at the protein level, we analyzed the secretion of chemokines using skin explants. CCL2, CCL3 and CXCL1 were more released upon papain treatment in the skin of mice deprived of dietary AhR ligands (Fig.5E), consistent with mRNA expression. Collectively, these results show that homeostatic activation of AhR via dietary ligands down-modulates inflammatory pathways in epidermal cells.

**Figure 5.**
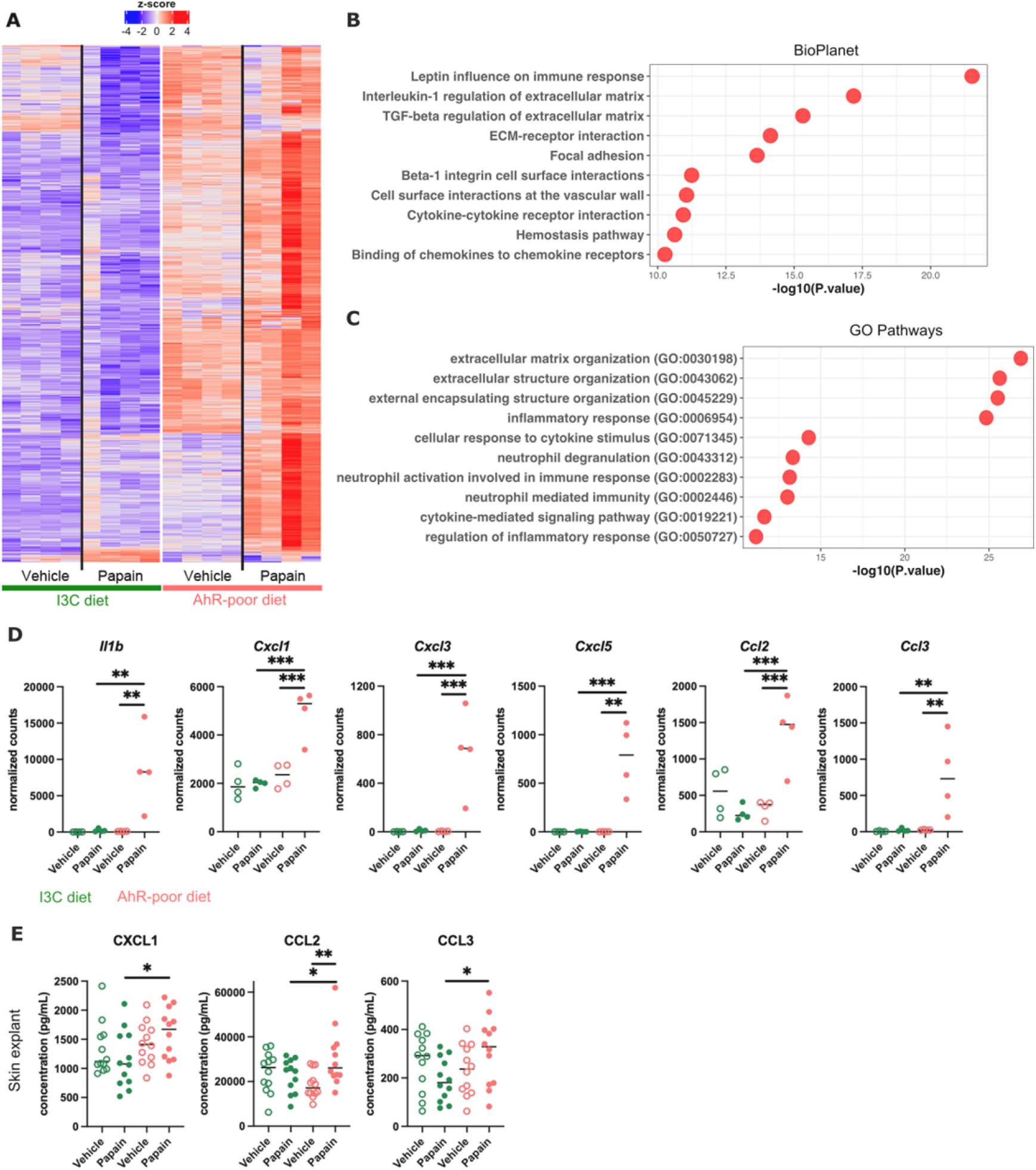
Dietary AhR ligands regulate the inflammatory profile of epidermal cells. Mice were fed on AhR-poor or I3C diet. Papain or vehicle (PBS) was injected in the footpad. (A-D) After 6h, epidermal cells were extracted and subjected to RNA-seq analysis. n=4 biological replicates. (A) Diet-dependent differentially expressed genes. (B-C) Enrichment for biological pathways using BioPlanet (B) or Gene Ontology Biological Process databases (C). (D) Normalized counts for selected genes, median is shown. One-way ANOVA. (E) After 24h, footpad skin was collected and explants were cultured for 24h to prepare conditioned medium. Chemokine concentrations were measured in the medium. Median is shown (n=12 in 3 independent experiments). Kruskal-Wallis test. For all panels * p<0.05; ** p<0.01; *** p<0.001.

### Microbiota-derived and diet-derived AhR ligands modulate different sets of epidermal genes

AhR is involved in keratinocyte differentiation and maintenance of skin barrier integrity (Haas et al., 2016; van den Bogaard et al., 2015). AhR ligands produced by commensal microbiota have been shown to be essential in this process (Uberoi et al., 2021). To address whether microbiota-derived and diet-derived AhR agonists have similar impact on epidermal genes, we re-analyzed public transcriptomic data from the epidermis of germ-free and Specific Pathogen-Free (SPF) mice (Uberoi et al., 2021). The GO signature for keratinocyte differentiation was enriched in the epidermis of SPF mice (Fig.6A), consistent with previous findings that microbiota-derived AhR ligands control the expression of genes involved in barrier function such as *Cdsn, Cldn1, Dsc1, Dsg1a, Flg, Ivl* and *Tjp3*, and of keratine molecules including *Krt10* (Uberoi et al., 2021) (Fig.6A). By contrast, the genes overexpressed in the epidermis of mice fed on the AhR-poor diet were not enriched in any group (Fig.6B). In particular, the expression of chemokine genes or *Itgb8* was comparable between SPF and germ-free mice (Fig.6B). In addition, we found that lack of dietary AhR ligands had no impact on, or even increased, the expression of keratinocyte barrier genes (Fig.6C). Consistent with this, we did not observe by histology any diet-dependent defect in *stratum corneum*, the outer layer of epidermis formed by cornified keratinocytes (Fig.1A and Fig.S1E). These observations suggest that the epidermal barrier function was not compromised in mice deprived of dietary AhR ligands and that AhR ligands derived from microbiota or diet modulate different set of genes in epidermal cells.

**Figure 6.**
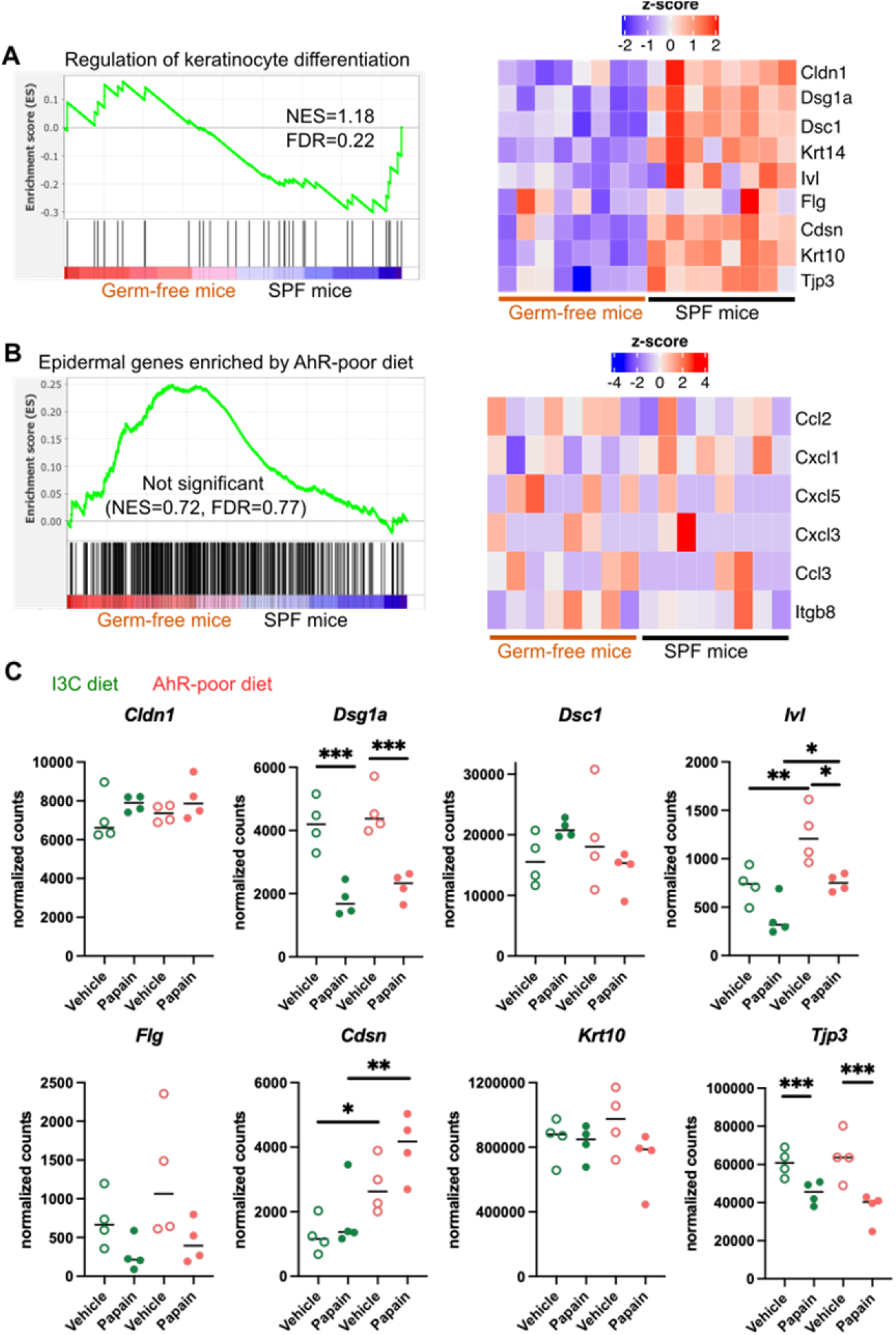
Microbiota-derived and diet-derived AhR ligands modulate different sets of epidermal genes. (A-B) RNA-seq data of the epidermis of germ-free or Specific Pathogen-Free (SPF) mice (n=8) was extracted from public source (GSE162925). Gene Set Enrichment Analysis was performed for Gene Ontology Biological Pathway ‘Regulation of keratinocyte differentiation’ signature (A) or the top 500 differentially expressed genes enriched in mice fed on AhR-poor diet (B). NES=normalized enrichment score. FDR=False Discovery Rate. Results are considered significant when NES>1 and FDR<0.25. (A-B) Scaled expression of selected genes. (C) Mice were fed on AhR-poor or I3C diet. Papain or vehicle (PBS) was injected in the footpad. After 6h, epidermal cells were extracted and subjected to RNA-seq analysis. n=4 biological replicates. Normalized counts for selected genes, median is shown. One-way ANOVA. For all panels * p<0.05; ** p<0.01; *** p<0.001; absence of star indicates ‘not significant’.

### AhR activation regulates keratinocyte production of TGF-β in mouse and human

Langerhans cells migration is regulated by Tgf-β, which retains Langerhans cells in the epidermis (Kel et al., 2010; Bobr et al., 2012; Mohammed et al., 2016). Bioactive Tgf-β can be produced from the latent form upon cleavage by integrin-β8 expressed on keratinocytes (Mohammed et al., 2016; Cambier et al., 2005). One of the pathways enriched in the epidermis of mice fed on the AhR-poor diet was related to Tgf-β regulation (Fig.5B). Indeed, expression of *Itgb8* was significantly higher in the epidermis of mice fed on the AhR-poor diet, with or without papain treatment (Fig.7A), suggesting increased release of bioactive Tgf-β in the skin of mice deprived of dietary AhR ligands. To directly test this, we measured total and bioactive Tgf-β release in skin explants. The concentration of bioactive Tgf-β, but not that of total Tgf-β, was significantly higher in the skin of mice fed on the AhR-poor diet (Fig.7B).

**Figure 7.**
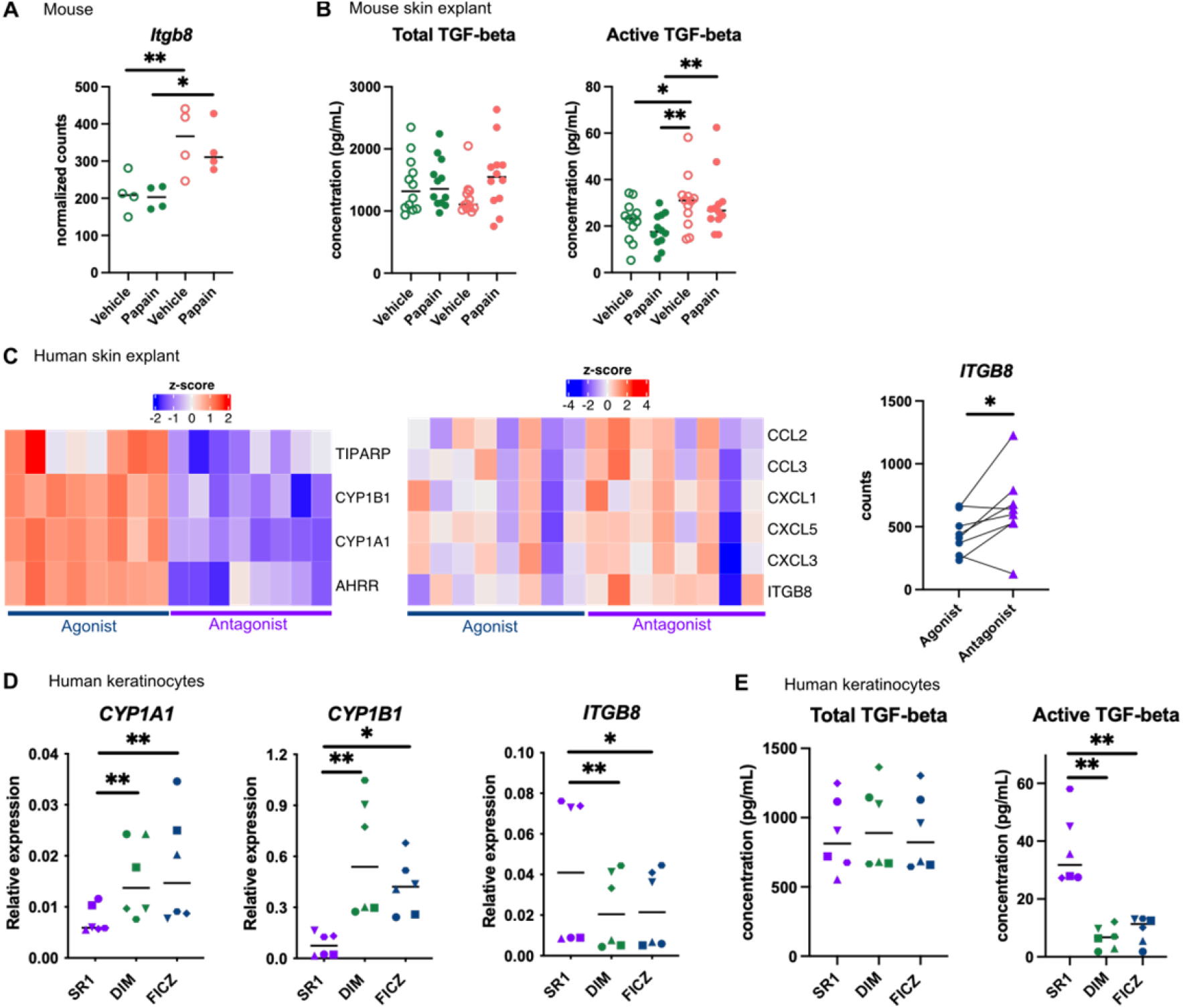
AhR activation regulates keratinocyte production of TGF-β in mouse and human. (A-B) Mice were fed on AhR-poor or I3C diet. Papain or vehicle (PBS) was injected in the footpad. (A) After 6h, epidermal cells were extracted and subjected to RNA-seq analysis. n=4 biological replicates. Normalized counts, median is shown. One-way ANOVA. (B) After 24h, footpad skin was collected and explants were cultured for 24h to prepare conditioned medium. Tgf-β concentration was measured in the medium. Median is shown (n=12 in 3 independent experiments). Kruskal-Wallis test. (C) RNA-seq data of human skin explant cultured in presence of AhR agonist (FICZ) or AhR antagonist (SR1) was extracted from public source (GSE47944) (n=8). Scaled expression of selected genes. Normalized counts for *ITGB8*, paired t-test. (D-E) Human HaCaT keratinocytes were cultured for 24h in presence of AhR antagonist (SR1) or AhR agonists (DIM or FICZ). (D) Expression of selected genes was measured by RT-qPCR. Median is shown (n=6 in 3 independent experiments, individual symbols represent paired conditions). Friedman test. (E) Tgf-β concentration was measured in the supernatant. Median is shown (n=6 in 3 independent experiments, individual symbols represent paired conditions). Friedman test. For all panels * p<0.05; ** p<0.01; *** p<0.001.

To address the relevance of these results in human, we first re-analyzed public transcriptomic data from human skin exposed *ex vivo* to an AhR antagonist (Stemregenin-1, SR1) or agonist (6-formylindolo(3,2-b)carbazole, FICZ) (Di Meglio et al., 2014). FICZ is an endogenous AhR ligand produced from the photo-oxidation of tryptophan (Öberg et al., 2005; Wincent et al., 2009). As expected, canonical AhR target genes were upregulated in the FICZ-treated samples (Fig7C). By contrast, chemokine genes (*CCL2, CCL3, CXCL1, CXCL3, CXCL5*) and *ITGB8* were more expressed in SR1-treated skin (Fig7C). These results are consistent with our transcriptomic data from mouse epidermis (Fig5D and Fig7A). To confirm that AhR activation regulates bioactive Tgf-β release by human keratinocytes, we used a human keratinocyte cell line (HaCaT cells). We cultured differentiated human keratinocytes in the presence of an AhR antagonist (SR1) or two different physiological AhR agonists, FICZ (produced by photo-oxidation) or DIM (produced from phytonutrients). AhR activation increased the expression of canonical target genes *CYP1A1* and *CYP1B1*, but decreased the expression of *ITGB8* (Fig7D). To directly address the impact of AhR activation on Tgf-β secretion, we measured total and bioactive Tgf-β in the supernatant. While total Tgf-β concentration was similar between treatments, bioactive Tgf-β was significantly more released by SR1-treated keratinocytes (Fig7E). Taken together, these results indicate that lack of AhR activation increases the production of bioactive Tgf-β in the skin in both mouse and human, thereby inhibiting Langerhans cells migration.

## Discussion

In this work, we showed that lack of dietary AhR ligands exacerbates cutaneous allergic Th2 responses and airway allergy after skin sensitization. We found that homeostatic activation of AhR by diet-derived agonists down-modulates inflammation pathways in epidermal cells. In particular, mice deprived of dietary AhR ligands displayed hyperproduction of bioactive Tgf-β in the skin, impairing Langerhans cells migration and their action as down-modulators of Th2 cells induction in the lymph nodes.

Previous work has reported a role for Langerhans cells in suppressing T cell responses in various models. In hapten-induced contact hypersensitivity, which is mediated by CD8 T cells, Langerhans cells are essential for tolerance to haptens (Kaplan et al., 2005), and to down-modulate T cell responses by producing IL10 (Igyarto et al., 2009) and by inducing regulatory CD4 T cells (Gomez de Agüero et al., 2012). In models of cutaneous sensitization to ovalbumin, Langerhans cells depletion increased Th2 cutaneous responses (Marschall et al., 2021; Luo et al., 2019) and T follicular helper cells (Marschall et al., 2021), as well as airway inflammation after intra-nasal recall (Marschall et al., 2021). Langerhans cells depletion also exacerbated lung Th2 responses after epicutaneous sensitization to house dust mite (Deckers et al., 2017b). It has been proposed that Langerhans cells produce IL10 upon exposure to allergens (Luo et al., 2019), but the mechanisms by which Langerhans cells regulate Th2 responses, and whether regulatory T cells are involved, remain to be better characterized. Consistent with these observations, we propose a model whereby dietary AhR ligands regulate the severity of cutaneous Th2 responses via the control of Langerhans cells migration, with decreased numbers of Langerhans cells in skin-draining lymph nodes leading to exacerbated Th2 cytokine production.

Previous studies have proposed a role for AhR in regulating airway inflammation. In a model of ovalbumin-induced airway allergy, AhR-deficient mice developed more severe allergic responses due to the increased ability of AhR-deficient T cells to proliferate and the higher activation of AhR-deficient lung DC (Thatcher et al., 2016). In addition, in another study using ovalbumin-induced airway allergy, systemic injection of high doses of the AhR agonist FICZ reduced eosinophilia and Th2 cytokines in the lung, and blood IgE levels (Jeong et al., 2012). By contrast, we found that airway allergic responses induced by papain exposure were not affected by the lack of dietary AhR ligands. Our results suggests that dietary AhR ligands do not play a role in the induction by lung DC of Th2 polarization *per se*,in cytokine secretion by lung ILC2 or in the inflammatory response of lung epithelial cells. This discrepancy could be explained by ligand-specific effects or by redundant effects of AhR agonists from different sources, the lack of dietary AhR ligand being compensated in the lung by endogenous or microbiota-derived ligands.

AhR can be activated by xenobiotics, diet-derived molecules, metabolites produced by microbiota metabolism, as well as endogenous ligands produced by cellular metabolism (Rothhammer and Quintana, 2019). Here we specifically focused on natural diet-derived AhR agonists, delivered by a physiological route, i.e food absorption. In epidermal cells, we found that dietary AhR agonists regulate a different set of genes than microbiota-derived AhR ligands. This is consistent with previous work identifying ligand-specific effects. In a model of MC903-induced atopic dermatitis, topical application or oral administration of microbiota-derived AhR ligand indole-3-aldehyde (IAId) reduced skin inflammation and type 2 responses, which was dependent on AhR (Yu et al., 2019). However, application of other AhR agonists such as kynurenin (an endogenous ligand) or indole-3-acetic acid (produced by microbiota) had no impact on disease symptoms (Yu et al., 2019). Mechanistically, it was shown that AhR activation by IAId regulated TSLP production by keratinocytes (Yu et al., 2019). By contrast, here we did not find any impact of dietary AhR ligands on TSLP production during papain-induced allergy. In a model of imiquimod-induced psoriasis, AhR-deficient mice had exacerbated skin inflammation (Di Meglio et al., 2014), while topical application of the AhR agonists FICZ (Di Meglio et al., 2014) or Tapinarof (Smith et al., 2017) ameliorated disease symptoms. Moreover, AhR-deficient keratinocytes were hyperresponsive to inflammatory stimuli *ex vivo* and overexpressed inflammatory mediators such as CXCL1 and CXCL5 (Di Meglio et al., 2014), consistent with our findings. Collectively, these observations highlight ligand-specific effects of AhR activation in keratinocytes.

In addition, intestinal type 3 ILC and lymphoid follicles are reduced in mice fed on an AhR-poor diet compared to I3C diet (Kiss et al., 2011), but are normal in germ-free mice (Lee et al., 2012), suggesting distinct effects of diet-derived and microbiota-derived AhR ligands on type 3 ILC and lymphoid follicles. The underlying mechanisms of such ligand-specific effects remain unclear (Rothhammer and Quintana, 2019).

In conclusion, we show that diet-derived ligands are a major source of homeostatic stimulation of AhR in keratinocytes, dampening their response to inflammation and regulating the release of bioactive Tgf-β, but not the barrier function. These results provide novel insight into the gut-skin axis, and could pave the way for optimizing diet interventions to reduce the development of cutaneous allergic diseases.

## Author Contributions

Investigation: AC, AdJ, RL, JLS, MM, SHC, ES. Conceptualization: ES. Formal analysis: AC, AdJ, JLS, MM, ES. Writing – original draft: ES. Writing – review and editing: all authors. Supervision: ES.

## Acknowledgements

This work was funded by INSERM, Institut Curie, Cancéropôle Ile-de-France, Institut National du Cancer (2018-1-PLBIO-01-ICR1) and Agence Nationale de la Recherche (ANR-10-LABX-0043, ANR-10-IDEX-0001-02 PSL, ANR-17-CE15-0011-01). The authors wish to thank the Flow Cytometry Core, the Pathex Platform, the NGS Platform, the Metabolomics and Lipidomics Platform and the In vivo Experiments Platform of Institut Curie. The authors thank S.Henri for helpful advice and M.Vocanson for providing founder Langerin-eGFP-DTR mice.

## Competing interests

The authors declare no competing interests.

## Material and Methods

### Animals

C57/B6J mice were obtained from Charles River (France). Mice were maintained on a purified diet (‘AhR-poor diet’, AIN-93M, Safe diets) supplemented or not in Indole-3-carbinol (I3C, 200 ppm, Sigma). In some experiments, mice were fed on a normal chow diet (4RF25 SV-PF 1609, Le comptoir des sciures). For Langerhans cells depletion, *Langerin-eGFP-DTR* mice (Kissenpfennig et al., 2005) were used with DTR^-/-^ littermates as controls, and injected intra-peritoneally with 500 ng of Diphteria Toxin (Sigma) 3 days prior to allergen treatment. For AhR *in vivo* inhibition, mice were treated by intra-peritoneal injection of 100 μg of AhR inhibitor CH-223191 (Invivogen) on 3 consecutive days prior to allergen treatment. Only female mice were used, except for the Langerhans cells depletion experiments. Mice were placed on the specific diet at 5 weeks of age for a period of adaptation of 3 weeks, before being used for any experiment. Mice were maintained under specific pathogen-free conditions at the animal facility of Institut Curie in accordance with institutional guidelines. Animal care and use for this study were performed in accordance with the recommendations of the European Community (2010/63/UE) for the care and use of laboratory animals. Experimental procedures were specifically approved by the ethics committee of the Institut Curie CEEA-IC #118 (Authorization APAFiS#24554-2020030818559195-v1 given by National Authority) in compliance with the international guidelines.

### Allergy models

For cutaneous allergy, mice were injected in the footpad with 50 μL of Phosphate Buffer Saline (PBS, vehicle) containing or not 50 μg of Papain (Sigma). Footpad skin was collected after 24h or 48h, popliteal lymph nodes were collected after 6 days. For airway allergy, mice were anesthetized using isofluorane and exposed intra-nasally to 20 μL of PBS (vehicle) containing or not 10 μg of Papain, at day 0, 1, 13 and 20. Blood was collected at day 20. Broncho-alveolar lavage, lungs and lymph nodes were collected at day 21.

For airway allergy after skin sensitization, mice were injected in the footpad with 50 μL of PBS (vehicle) containing or not 50 μg of Papain (Sigma). At day 7, all mice were placed on the I3C diet. Mice were exposed intra-nasally to 20 μL of PBS (vehicle) containing or not 10 μg of Papain at day 14, 15, 27 and 35. Bronchoalveolar lavage and lungs were collected at day 36.

### IgE measurement

Blood was collected and left at room temperature for 3h to coagulate. After centrifugation (450g, 10 min), serum was collected for analysis and kept at −20°C. IgE concentration was measured using ELISA (Invitrogen). The limit of detection was 140 pg/mL.

### Histology

Whole feet from treated mice were collected and fixed in Formalin. Samples were decalcified in RDO (Eurobio scientific) for 6h at 37°C before paraffin embedding. Samples were cut into 3 μm thin sections, deparaffinized and stained with Hematoxylin (Dako), Eosin (RAL diagnostics) and Safran (RAL diagnosctics) (HES). Slides were scanned with Philips ULTRA FAST scanner 1.6 RA. Images were analyzed using *QuPath* (v.0.3.1) (Bankhead et al., 2017).

### Indole measurement in mouse blood

Blood samples were collected in EDTA coated tubes. Samples were centrifuged for 15 min at 1200g to obtain serum. Indoles levels were measured in serum using liquid chromatography coupled to high-resolution mass spectrometry (HPLC-HRMS) (Lefèvre et al., 2019). Briefly, 50-100 μL of serum were added to 800μL of methanol and samples were vortexed for 5 min in a thermomixer at 4°C followed by an incubation at −20 °C for 30 min. After centrifugation at 13,300g for 10 min, the supernatant was collected and concentrated using a SpeedVac vacuum concentrator (Thermo Scientific). Samples were resuspended in 100 μL of 10% methanol solution, vortexed in a thermomixer for 5 min and centrifuged at 13,300g for 10 min. 80 μL of each sample was transferred to a liquid chromatography vial and 10 μL injected in the HPLC-HRMS system. Chromatography was carried out with a Phenomenex Kinetex 1.7 μm XB – C18 (150 mm × 2.10 mm) and 100 Å HPLC column maintained at 55 °C. The solvent system comprised mobile phase A [0.5% (vol/vol) formic acid in water], and mobile phase B [0.5% (vol/vol) formic acid in methanol]. The gradient was set-up as follows: 0-2 min, 0% B; 2-7 min, 0-50% B; 7-15 min, 50-100% B; 15-18 min, 100% B, 18-18.5 min 100-0% B and 18.5-21.5 min 0%B. HRMS analyses were performed on a HPLC Vantage Flex (Thermo Fischer Scientific) coupled to a Q-Exactive Focus mass spectrometer (Thermo Fisher Scientific) that was operated in positive (ESI+). The HPLC autosampler temperature was set at 4°C. The heated electrospray ionization source was set with a spray voltage of 4.5 kV, a capillary temperature of 250°C, a heater temperature of 475°C, a sheath gas flow of 35 arbitrary units (AU), an auxiliary gas flow of 10 AU, a spare gas flow of 1 AU, and a tube lens voltage of 100 V. During the HRMS acquisition, the scan range was set to m/z=100-500 Da, the instrument operated at 70,000 resolution (m/z = 200), with an automatic gain control (AGC) target of 1×10^6^ and a maximum injection time (IT) set to automatic. Instrumental chromatography stability was evaluated by injection of a synthetic standard mixture with all metabolites of interest and quality control (QC) samples quality control obtained from a pool of the left-over of all samples analyzed. This QC sample was reinjected once at the beginning of the analysis, every 10 sample injections, and at the end of the run. Ionization and retention times were validated with pure standards and are summarized in the following table:

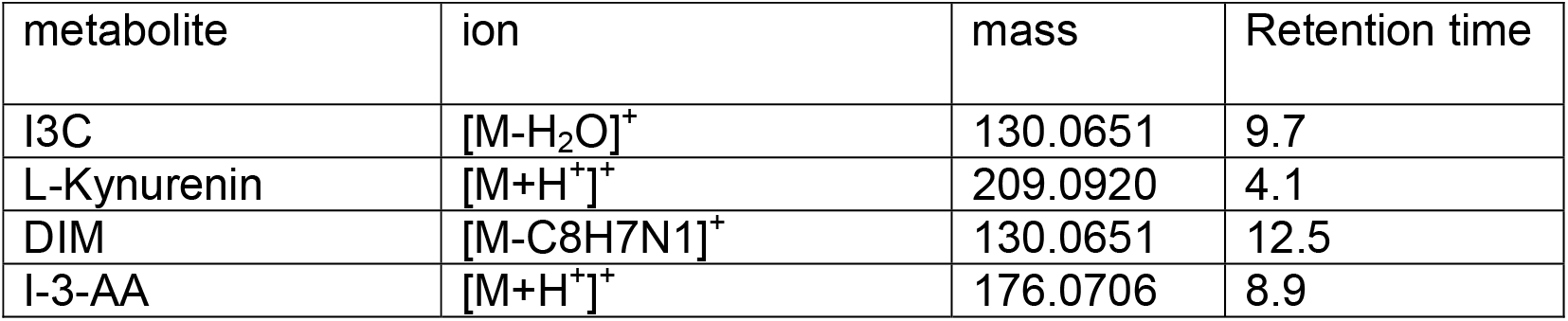

| metabolite | ion | mass | Retention time |
| --- | --- | --- | --- |
| I3C | $[M-H_2O]^+$ | 130.0651 | 9.7 |
| L-Kynurenin | $[M+H^+]^+$ | 209.0920 | 4.1 |
| DIM | $[M-C_8H_7N_1]^+$ | 130.0651 | 12.5 |
| I-3-AA | $[M+H^+]^+$ | 176.0706 | 8.9 |

Peak area was used as read out for relative quantification of the metabolites across the samples and normalized by serum volume.

### Skin explant analysis

Skin from the footpad was harvested 24h after vehicle or papain treatment. Skin explants from both footpads were pooled and incubated for 24h in 1 mL of RPMI (Gibco) medium containing 10% fetal calf serum (FCS, Biosera). After centrifugation (450g, 5 min), conditioned medium was collected for analysis and kept at −20°C. Tgf-β concentration was measured using ELISA (Total Tgf-β1 Legend MAX and Free Active Tgf-β1 Legend MAX, Biolegend). The limit of detection was 8 pg/mL for Tgf-β1. CCL2, CCL3 and CXCL1 concentration was measured using CBA (BD Biosciences). The limit of detection was 10 pg/mL.

### Flow cytometry

Cells were stained with indicated antibody cocktails supplemented with Fc block (BD Biosciences) in FACS buffer (PBS containing 0.5% BSA and 2mM EDTA) for 30-45 min on ice. After washing with FACS buffer, cells were resuspended in FACS buffer containing DAPI (Fischer Scientific, 100ng/mL). Cells were acquired on a ZE5 (Bio-Rad) or FACSVerse (BD Biosciences) instrument. Supervised analysis was performed using FlowJo software v10 (FlowJo LLC).

### Broncho-alveolar lavage analysis

Broncho-alveolar lavage was collected by injection of 1 mL of PBS in the bronchoalveolar space. Suspensions were filtered using 40μm cell strainers. After centrifugation (450g, 5 min), the lavage was collected for analysis of soluble mediators and kept at −20°C. IL5 and IL13 concentration was measured using Enhanced Sensitivity CBA (BD Biosciences). The limit of detection was 274 fg/mL. Cells were stained with anti-TCRβ BUV737 (BD Biosciences, clone H57-597), anti-CD11c PerCpCy5.5 (BD Biosciences, clone HL3), anti-SiglecF BV480 (BD Biosciences, clone E50-2440), anti-CD45 FITC (BioLegend, clone 30-F11), anti-CD11b Pe-CF594 (BD Biosciences, clone M1/70), anti-MHCII BV786 (BioLegend, clone M5/114.15.2), anti-Ly6G BV605 (BioLegend, clone 1A8), anti-CD4 BV650 (Biolegend, clone RM4-5), anti-Ly6C AF700 (BioLegend, clone HK1.4).

### Lymph node cells analysis

For Th2 polarization analysis, lymph nodes were collected and dissociated by forcing through a 40μm cell strainer. Cell suspensions were analyzed by flow cytometry after staining with anti-TCRβ APC (BioLegend, clone H57-597), anti-CD11b PerCpCy5.5 (BD Biosciences, clone M1/70), anti-CD4 FITC (BD Biosciences, clone RM4-5) and anti-CD19 APC-Cy7 (BD Biosciences, clone 1D3). Normalized cell numbers (2×10^5^ cells/well) were cultured for 24h in 100 μL of RPMI medium containing 10% FCS in the presence of 5 μL of anti-CD3/CD28 beads (ThermoFisher). After centrifugation (450g, 5 min), supernatant was collected for analysis and kept at −20°C. IL4, IL5, IL10, IFN-γ, IL17A and IL13 concentration was measured using Enhanced Sensitivity CBA (BD Biosciences). The limit of detection was 274 fg/mL.

For flow cytometry of DC, lymph nodes were cut into small pieces and incubated for 30 minutes at 37°C in digestion mix: RPMI containing 0.5mg/mL DNAse I (Sigma) and 0.5mg/mL collagenase D (Roche). Cell suspensions were then filtered using 40μm cell strainers. Antibodies used were anti-CD172a BUV737 (BD Biosciences, clone P84), anti-CD19 BV480 (BD Biosciences, clone 1D3), anti-CD3 BV480 (BD Biosciences, clone 500A2), anti-XCR1 BV510 (Biolegend, clone ZET), anti-CD11c BV785 (Biolegend, clone N418), anti-CD86 FITC (BD Biosciences, clone GL1), anti-CD26 PE (Biolegend, clone H194-112), anti-CD40 PerCP-efluor710 (eBioscience, clone 1C10), anti-PDL2 APC (Biolegend, clone TY25), anti-MHC II BV650 (Biolegend, clone M5/114.15.2), anti-EpCAM APCFire750 (BioLegend, clone G8.8).

### Epidermal cells analysis

Skin from the footpad was harvested using scalpels. The epidermis and dermis layers were separated after incubation at 37°C for 1h in 0.4 mg/mL dispase II (Roche). The epidermis was collected and then cut into small pieces using scalpels and incubated for 30 minutes with agitation at 37°C in RPMI containing 10% FCS and 0.5 mg/mL DNAse I. Suspensions were filtered using 40μm cell strainers.

For Langerhans cells analysis, after centrifugation (450g, 5 min) cells were stained with anti-CD45 FITC (BioLegend, clone 30-F11), anti-EpCAM APCFire750 (BioLegend, clone G8.8) and anti-CD11b PerCPCy5.5 (BD Biosciences, clone M1/70).

For RNA-seq analysis, after centrifugation (150g, 5 min), dead cells were removed using EasySep Dead cell removal kit (StemCell). After this step, viability was around 70% as assessed by flow cytometry. Epidermal cells were composed of 95% keratinocytes (CD45-cells) as assessed by flow cytometry.

### Imaging of epidermis

For imaging*, Langerin*-eGFP-DTR mice were used. Mice were placed on the AhR-poor or I3C diet for 3 weeks. Epidermis was prepared from ear skin after hair removal using a depilating cream (Veet). The epidermis and dermis layers were separated after incubation at 4°C for 16h in 0.2 mg/mL dispase II (Roche). Epidermal layers were fixed in 4% Para-formaldehyde for 20 min at room temperature. After washing in PBS, epidermal layers were placed on coverslips and preserved using Fluoromount-G mounting medium (SouthernBiotech).

eGFP fluorescence was imaged on an inverted laser scanning confocal microscope (Leica DMI8 with a sp8 scanning unit) equipped with an oil immersion objective (40x, NA=1.35). The 488 nm laser was used for excitation and eGFP signal was collected on an Hybrid detector. A pixel size of 0.28 μm was chosen and Z stacks of 3 planes were acquired (Z step = 1 micron).

Image analysis was performed using Fiji software (Schindelin et al., 2012). To analyze Langerin+ cells density, a homemade macro was used. After projection of the 3 planes, a mask of the total eGFP was obtained. In a second step, images were blurred using a large radius (3.4 microns) to distinguish cellular stroma and count the number of cells. Finally, the number of cells was normalized to the tissue area to compute cell density.

### RNA-seq library preparation

Epidermal cells were isolated 6h after vehicle or Papain treatment. Cells were lysed in RLT buffer (QIAGEN). Total RNA was extracted using the RNAeasy minikit (Qiagen) including on-column DNase digestion according to the manufacturer’s protocol. The integrity of the RNA was confirmed in BioAnalyzer using RNA 6000 Nano kit (Agilent Technologies) (8.8 < RIN < 10). Libraries were prepared according to Illumina’s instructions accompanying the TruSeq Stranded mRNA Library Prep Kit (Illumina). 500 ng of RNA was used for each sample. Library length profiles were controlled with the LabChip GXTouchHT system (Perkin Elmer). Sequencing was performed in 3 sequencing unit of NovaSeq 6000 (Illumina) (100-nt-length reads, paired end) with an average depth of 40 millions of clusters per sample.

### RNA-seq data analysis

Genome assembly was based on the Genome Reference Consortium (mm10). Quality of RNA-seq data was assessed using *FastQC*. Reads were aligned to the transcriptome using *STAR* (Dobin et al., 2013). Differential gene expression analysis was performed using *DESeq2* (v1.22.2)(Love et al., 2014). Genes with low number of counts (<10) were filtered out. Differentially expressed genes between ‘Vehicle’ and ‘Papain’ treatment for each diet, or between ‘AhR-poor diet’ and ‘I3C diet’ conditions for each treatment, were calculated using the design ‘group’. Differentially expressed genes were identified based on adjusted p-value < 0.01 and Log2 FoldChange > 1. Complete gene lists are included in Table S1. Heatmaps of log2-scaled expression were generated with *ComplexHeatmap*. Pathway enrichment was analyzed using *EnrichR* (Kuleshov et al., 2016). Sequencing data has been deposited in GEO (accession number GSE198368).

### Analysis of public transcriptomic data

Data was downloaded from GEO. Raw count matrix was normalized using *DESeq2*.For human skin explant (GSE47944) (Di Meglio et al., 2014), data from non-lesional skin exposed to SR1 or FICZ was used. For mouse epidermis (GSE162925) (Uberoi et al., 2021), data from germ-free and Specific Pathogen-free mice were used. Heatmaps of log2-scaled expression were generated with *ComplexHeatmap*.

### Gene Set Enrichment Analysis

GSEA (Subramanian et al., 2005) was performed using the GSEA software (version 4.0.3) with the default parameters, except for the number of permutations that we fixed at n=1000. Results are considered significant when NES>1 and FDR<0.25. RNA-seq data from the epidermis of germ-free and Specific Pathogen-free mice (GSE162925) was normalized using *DESeq2*. Gene signature for keratinocyte differentiation (GO:0045616) was downloaded from MSigDB (v7.5.1) (Liberzon et al., 2015). Gene signature for epidermal cells in AhR-poor diet was obtained by using the top 500 differentially expressed genes upregulated in ‘AhR-poor diet’ versus ‘I3C diet’ for vehicle treatment, based on log2 fold change.

### Human keratinocyte culture

Human HaCaT keratinocytes were cultured with DMEM medium without glutamine or calcium (Gibco) supplemented with antibiotics (penicillin and streptomicin), 10% fetal calf serum (FCS) treated with Chelex (Sigma) to removed endogenous calcium, and 0.03 mM calcium chloride (low calcium growth medium). Cells were kept at 80% confluence in low calcium growth medium, in order to keep a basal undifferentiated phenotype (Wilson, 2014). For differentiation, cells were switched to the same medium containing 2.8 mM calcium chloride (high calcium growth medium).

HaCaT cells were exposed for 24h to 8μM SR1 (Cayman chemicals), 5 μg/mL Diindolylmethane (DIM, Sigma) or 60 nM 6-Formylindolo[3,2-b]carbazole (FICZ, Enzo Life Sciences) in low calcium growth medium. Medium was then replaced by high calcium growth medium and cells were further cultured for 24h. Cells were lysed in RLT buffer and supernatants were collected for analysis using ELISA (Total Tgf-β1 Legend MAX and Free Active Tgf-β1 Legend MAX, Biolegend). The limit of detection was 8 pg/mL for Tgf-β1.

### qPCR

Cells were harvested and lysed in RLT buffer (QIAGEN). RNA extraction was carried out using the RNAeasy micro kit (QIAGEN) according to manufacturer’s instructions. Total RNA was retro-transcribed using the superscript II polymerase (Invitrogen), in combination with random hexamers, oligo dT and dNTPs (Promega). Transcripts were quantified by real time PCR on a 480 LightCycler instrument (Roche). Reactions were carried out in 10μL, using a master mix (Eurogentec), with the following Taqman Assays primers (Merck): *Cyp1a1* (Mm00487218_m1), *Muc5ac* (Mm01276705_g1), *Gob5* (Mm01320697_m1), *Gapdh* (Mm99999915_g1), *B2m* (Mm00437762_m1), *Polr2a* (Mm00839502_m1), *CYP1B1* (Hs00164383_m1), *CYP1A1* (Hs01054796_g1), *ITGB8* (Hs00174456_m1), *B2M* (HS00187842_m1), *HPRT* (Hs02800695_m1), *RPL34* (Hs00241560_m1). The second derivative method was used to determine each Cp and the expression of genes of interest relative to the housekeeping genes (*Gapdh, B2m, Polr2a* for mouse and *B2M, HPRT, RPL34* for human) was quantified.

### Statistical analysis

Statistical tests were performed using Prism v9 (GraphPad Software). Statistical details for each experiment can be found in the corresponding figure legend. N corresponds to the number of biological replicates. Absence of star indicates ‘non-significant’. ANOVA was performed with Tukey’s multiple comparisons test.

## Supplementary material

**Figure S1.**
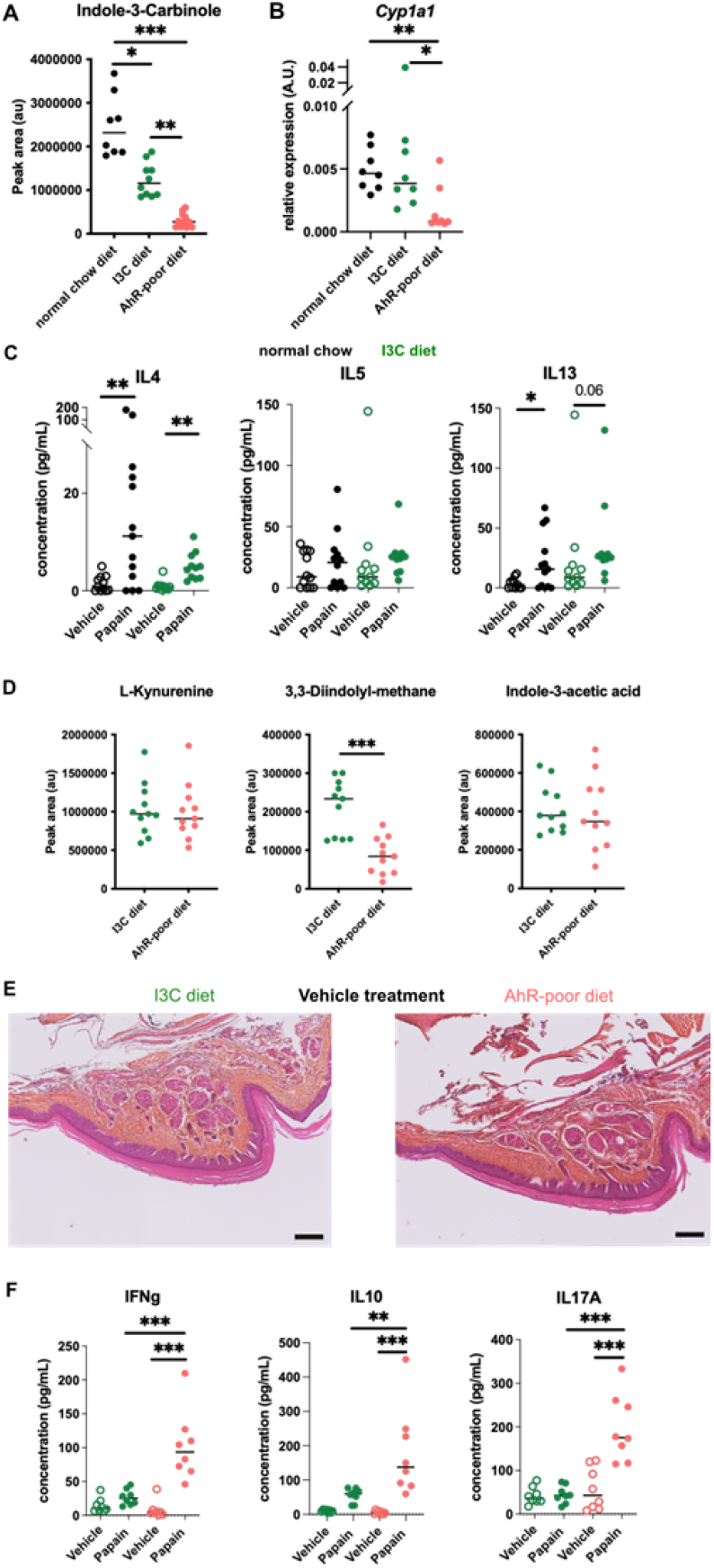
Comparable cutaneous allergic Th2 responses in mice fed with normal chow and I3C diet. Mice were fed on a normal chow or a AhR-poor diet or enriched in Indole-3-carbinol (I3C diet). (A) Relative abundance of I3C was measured in the serum. Median is shown (n=8-11 in 2 independent experiments). (B) Liver lysates were analyzed by RT-qPCR. Median is shown (n=8 in 2 independent experiments). One-way ANOVA. (C) Papain or vehicle (PBS) was injected in the footpad. After 6 days, normalized numbers of lymph node T cells were restimulated ex vivo and cytokine secretion was measured in the supernatant. Median is shown (n=11-13 in 3 independent experiments). Kruskal-Wallis test. (D) Relative abundance of AhR agonists was measured in the serum. Median is shown (n=11 in 2 independent experiments). (E) 24h after vehicle injection, tissues were analyzed by histology (Hematoxylin, Eosin and Safran staining). Representative results (n=6 per condition). Bar = 100 μm. (F) After 6 days, normalized numbers of lymph node T cells were restimulated ex vivo and cytokine secretion was measured in the supernatant. Median is shown (n=11-12 in 3 independent experiments). Kruskal-Wallis test. For all panels * p<0.05; ** p<0.01; *** p<0.001.

**Figure S2.**
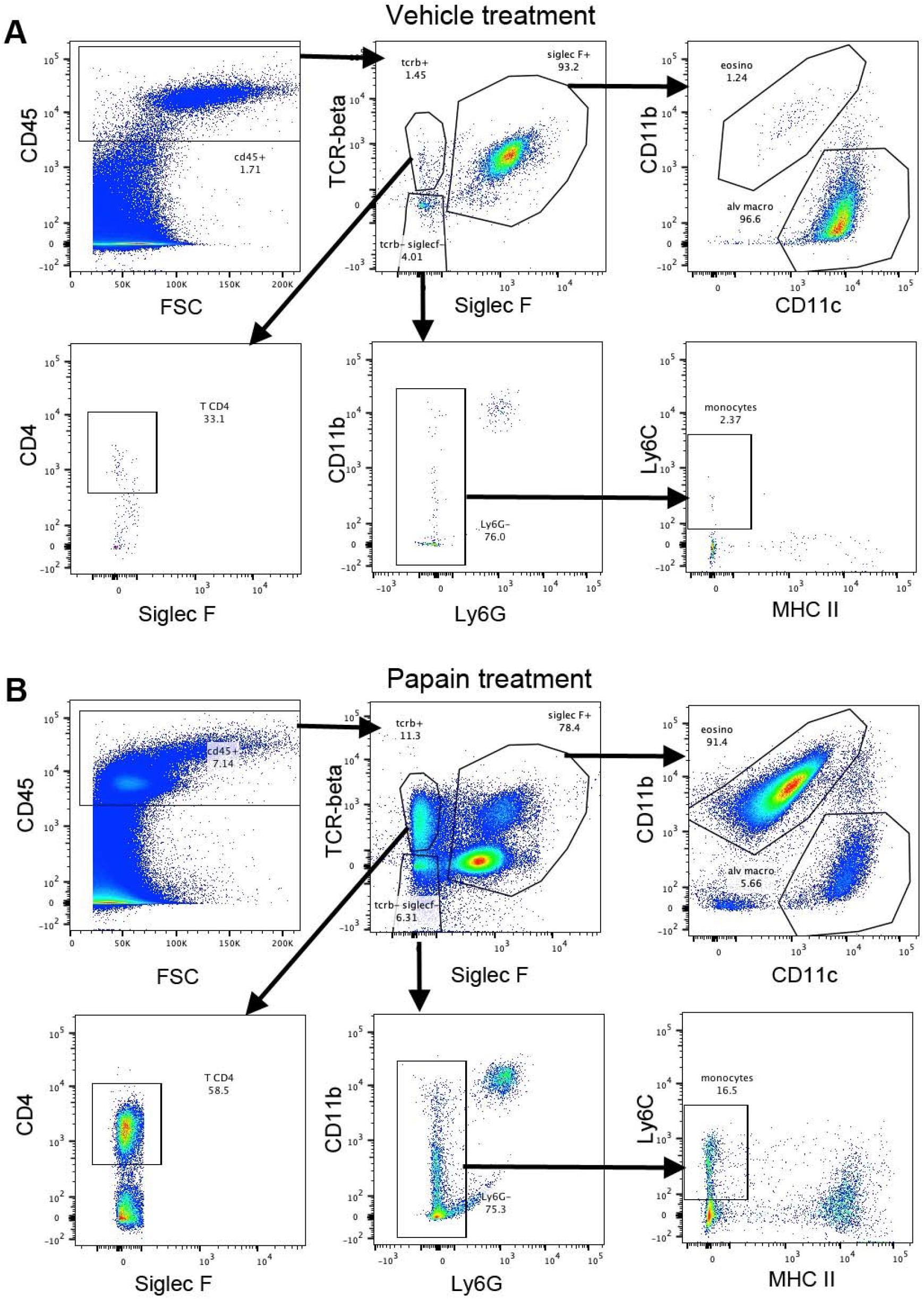
Gating strategy for broncho-alveolar cells. Broncho-alveolar cells were gated on live singlets. One representative example for mice treated with vehicle (A) or papain (B).

**Figure S3.**
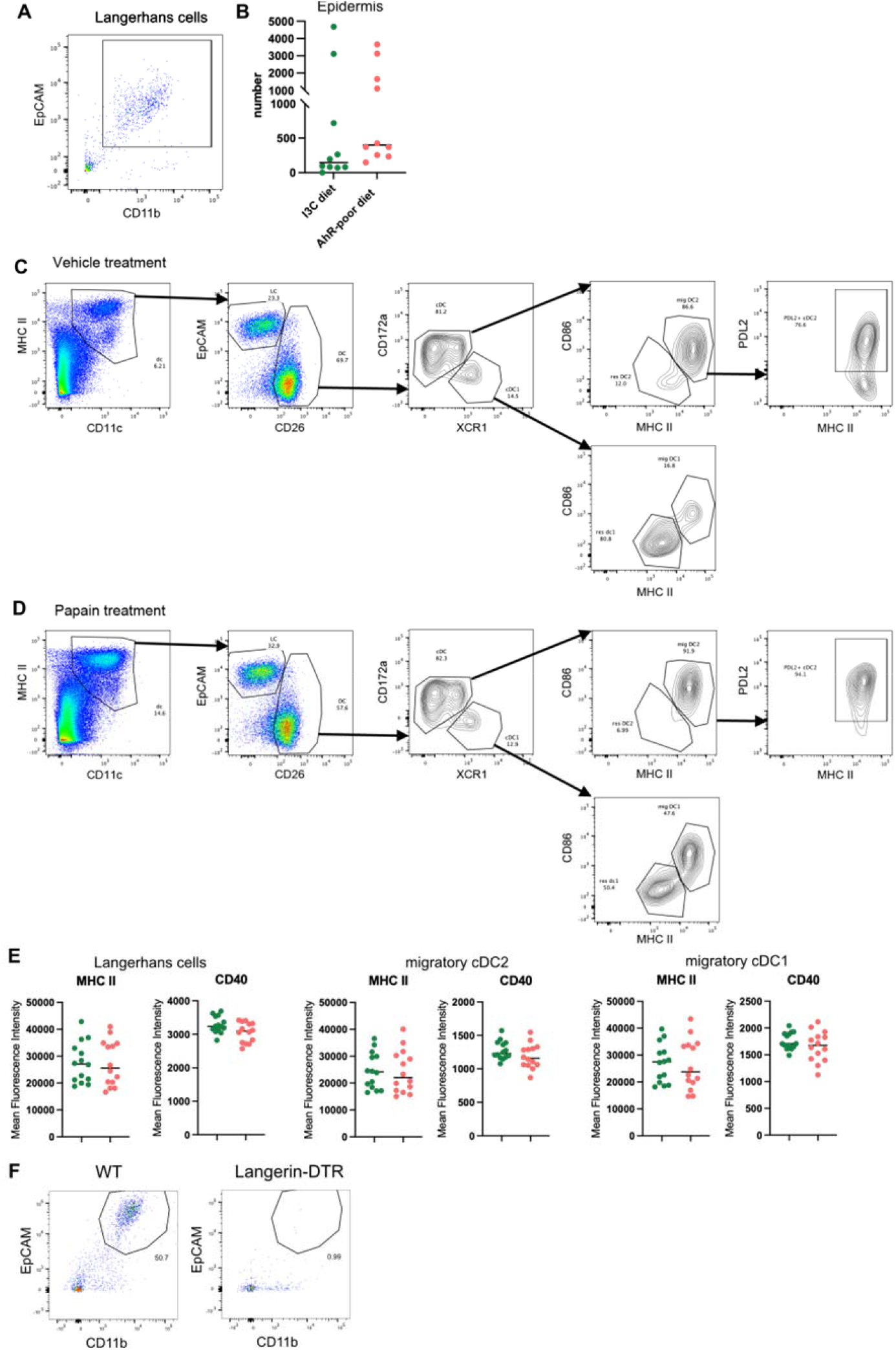
Analysis of cutaneous allergy. (A) Gating strategy for epidermal Langerhans cells. Cells were gated on live singlets CD45^+^ cells. (B) Mice were fed on AhR-poor or I3C diet. Number of Langerhans cells in the footpad epidermis. Median is shown (n=10 in 3 independent experiments). (C-D) Gating strategy for popliteal lymph nodes analysis. Lymph node cells were gated on live singlets CD19^-^ CD3^-^cells. One representative example for mice treated with vehicle (C) or papain (D). (E) Mice were fed on AhR-poor or I3C diet. Mean Fluorescence intensity for MHC II molecules and CD40 in indicated cell types 48h after papain injection. Median is shown (n=14 in 3 independent experiments). (F) Langerin-DTR mice and WT littermates were injected with Diphteria Toxin. After 3 days, footpad epidermis was analyzed by flow cytometry. Representative results (n=5 in 2 independent experiments). For all panels, absence of star indicates ‘not significant’.

**Figure S4.**
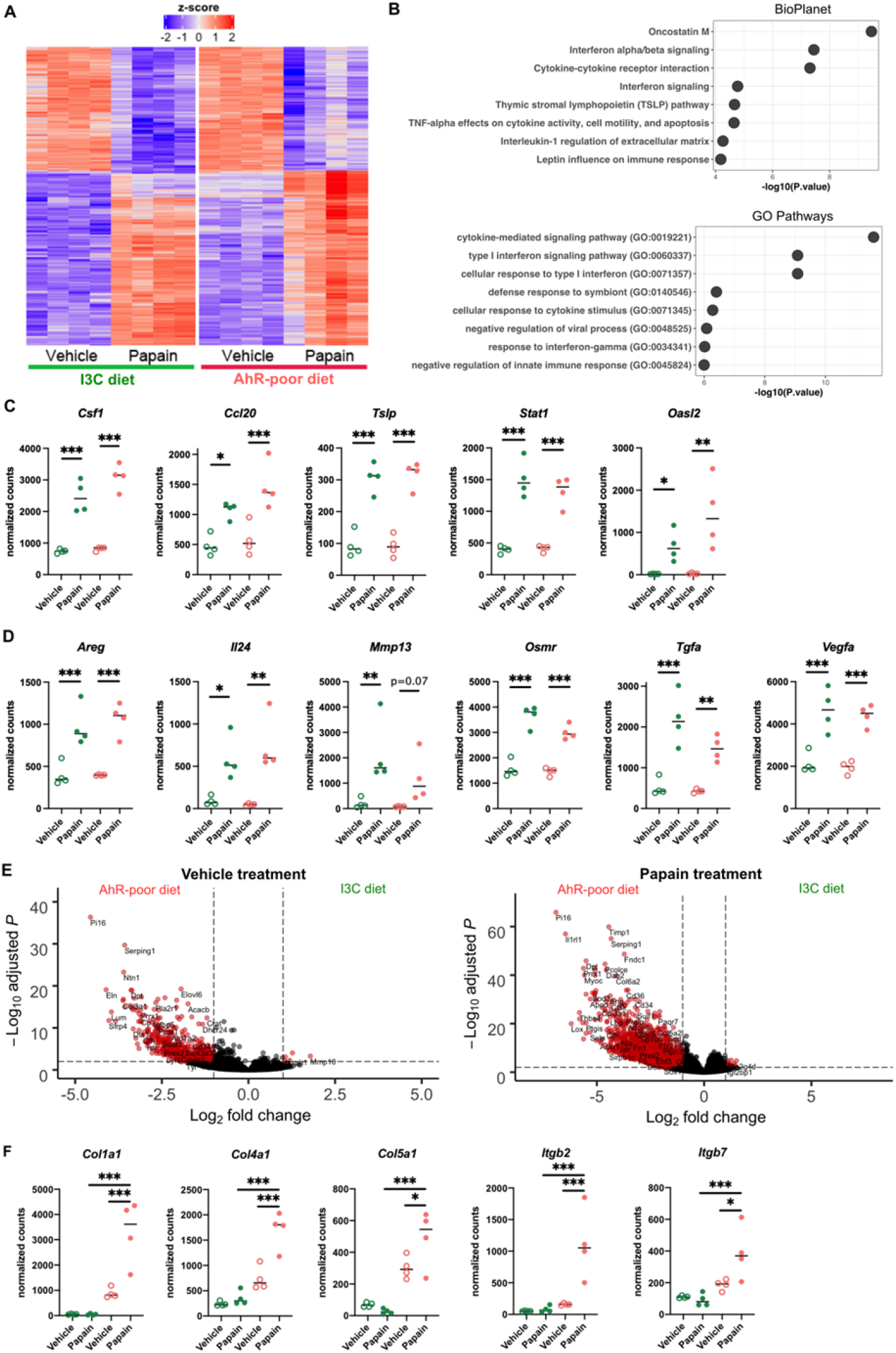
Dietary AhR ligands modulate the transcriptomic profile of epidermal cells. Mice were fed on AhR-poor or I3C diet. Papain or vehicle (PBS) was injected in the footpad. After 6h, epidermal cells were extracted and subjected to RNA-seq analysis. n=4 biological replicates. (A) Diet-independent differentially expressed genes. (B) Enrichment for biological pathways using BioPlanet or Gene Ontology Biological Process databases. (C-D) Normalized counts for selected genes, median is shown. One-way ANOVA. (E) Differentially expressed genes between mice fed on AhR-poor or I3C diet in Vehicle-treated mice (left) or Papain-treated mice (right). (F) Normalized counts for selected genes, median is shown. One-way ANOVA. For all panels * p<0.05; ** p<0.01; *** p<0.001.

**Table S1. Lists of differentially expressed genes**. Differentially expressed genes between ‘Vehicle’ and ‘Papain’ treatment for each diet, or between ‘AhR-poor diet’ and ‘I3C diet’ conditions for each treatment. Adjusted p-value and Log2 FoldChange are indicated.

